# Dissociating endogenous and exogenous delta activity during natural speech comprehension

**DOI:** 10.1101/2024.02.01.578181

**Authors:** Nikos Chalas, Lars Meyer, Chia-Wen Lo, Hyojin Park, Daniel S. Kluger, Omid Abbasi, Christoph Kayser, Robert Nitsch, Joachim Gross

**Affiliations:** Institute for Biomagnetism and Biosignal Analysis, University of Münster, Münster, Germany; Otto-Creutzfeldt-Center for Cognitive and Behavioral Neuroscience, University of Münster, Münster, Germany; Max Planck Institute for Human Cognitive and Brain Sciences, Leipzig, Germany; Centre for Human Brain Health (CHBH), School of Psychology, University of Birmingham, Birmingham, United Kingdom; Department for Cognitive Neuroscience, Faculty of Biology, Bielefeld University, 33615 Bielefeld, Germany; Institute for Translational Neuroscience, University of Münster, Münster, Germany

**Author notes:** Correspondence: Nikos Chalas, University of Münster, Institute for Biomagnetism and Biosignal Analysis, Malmedyweg 15, 48149 Münster, Germany;.

## Abstract

Decoding human speech requires the brain to segment the incoming acoustic signal into meaningful linguistic units, ranging from syllables and words to phrases. Integrating these linguistic constituents into a coherent percept sets the root of compositional meaning and hence understanding. One important cue for segmentation in natural speech are prosodic cues, such as pauses, but their interplay with higher-level linguistic processing is still unknown. Here we dissociate the neural tracking of prosodic pauses from the segmentation of multi-word chunks using magnetoencephalography (MEG). We find that manipulating the regularity of pauses disrupts slow speech-brain tracking bilaterally in auditory areas (below 2 Hz) and in turn increases left-lateralized coherence of higher frequency auditory activity at speech onsets (around 25 - 45 Hz). Critically, we also find that multi-word chunks—defined as short, coherent bundles of inter-word dependencies—are processed through the rhythmic fluctuations of low frequency activity (below 2 Hz) bilaterally and independently of prosodic cues. Importantly, low-frequency alignment at chunk onsets increases the accuracy of an encoding model in bilateral auditory and frontal areas, while controlling for the effect of acoustics. Our findings provide novel insights into the neural basis of speech perception, demonstrating that both acoustic features (prosodic cues) and abstract processing at the multi-word timescale are underpinned independently by low-frequency electrophysiological brain activity.

## Introduction

Natural speech comprises a rich spectrotemporal signal [1,2] from which, after a set of computations [3,4], we gain access to language comprehension. Over the last decades a plethora of studies investigated associations between brain activity and continuous speech during listening. One significant finding suggests that acoustical sampling is implemented via temporally aligning windows of high excitability, orchestrated by rhythmic brain activity at multiple time scales to relevant parts of the stimulus [5–11]. In this line, it still remains unclear whether parsing auditory speech through rhythmical activity depends only on exogenous speech rhythms or also on endogenous, top-down rhythms.

For speech, syllabicity occurs on a timescale of ∼200 ms [12,13], which aligns with the rhythmicity in the amplitude modulations of speech (around 5 Hz; [14]) and is hypothesized to be entrained - and therefore be tracked - by theta-band oscillator at a similar timescale (4 - 7 Hz [8,15–17]). However, on the timescale of word groups such an alignment between acoustic components, linguistic elements and brain activity is not clearly evident. One prominent example are prosodic phrases, which exhibit rhythmicity in the scale of ∼1 s [18,19] and are proposed to be tracked accordingly by an acoustically-driven oscillator (in the delta range; below 2 Hz; [20–22]). Acoustic – and prosodic – landmarks, speech pauses, have been also related to speech-brain alignment in the delta frequency band [23].

Previously, neural tracking of prosody has been reported in the delta timescale [24]. Yet, factorial experiments suggest that delta-band tracking of prosody may in part be confounded by neural activity that relates to the segmentation and content-level integration of abstract multi-word units. For example, the phase of ongoing delta activity was found to predict termination of multi-word chunks [25] by endogenously establishing a time-constraint for segmentation in the absence of prosodic cues [26]. The most influential evidence of delta activity in the processing of multi-word chunks was reported in an artificially constructed paradigm in which isochronous syllables were combined hierarchically into phrases and sentences [27]. Spectral peaks in the brain signal matching the timescale of the phrasal units were considered as an indication of internal and contextually-dependent processing. Similar findings were reported by incorporating more sophisticated paradigms [28,29]. Yet, again, this line of research has also been criticized to confound effects stemming, at least partially, from non-syntactic (implicit prosodic) factors [30,31]. In fact, syntax and prosody are ontologically [32] and temporally [33] intertwined. It is thus not surprising that prosodic boundaries facilitate higher-level syntactic processing [34–36], and more prominently so in the early-stage of language acquisition [37]. Given that in defining phrasal boundaries, bottom-up acoustical properties and top-down combinatorial processing are difficult to disentangle, it is unclear whether the attributed effect of syntactic processing echoes prosodic properties of the input (e.g. pauses or pitch contour), evident in low-frequency (below 2 Hz) brain activity. Importantly, top-down entrainment in the delta range has been found to occur for auditory rhythmic tone patterns, lacking physical boundaries [38]. But so far, systematic research investigating endogenous delta rhythms with respect to higher linguistic processing during natural speech comprehension is lacking.

Here we aim to address a fundamental mechanism of human speech processing: namely, how bottom-up exogenous and top-down endogenous combinatorial processing interact and shape perception during speech comprehension. For this we aimed to disentangle the sensory processing of prosody from the segmentation and integration of multi-word chunks. We temporarily altered prosodic boundaries in a naturalistic narrative (speech pauses; [39]); concurrently, we defined multi-word chunks as coherent bundles of inter-word dependencies (e.g., dependencies between a determiner, adjective, and a noun), using an independent computational formalization based on dependency annotations [40–44] We provide evidence that in spoken narratives both acoustic cues and abstract multi-word chunks are encoded separately in the phase of delta band auditory activity.

## Results

### Prosodic delta speech tracking and gamma coherence are anti-correlated during listening

We aimed to disentangle sensory- from contextually-driven encoding of speech with a focus on the phrasal timescale. To this end, N = 30 healthy participants listened to a story in two parts while their brain activity was recorded non-invasively with magnetoencephalography (MEG). First, we focused on altering the distributional statistics of speech, which we anticipated to lead to misalignment of speech-brain in the delta range. Thus we first asked how the acoustic natural statistics of connected speech drive sensory-cortical processing. For this, we divided the story into two equal blocks: In the first block the story was left intact (we will refer to it as *control* condition) whereas in the other, we identified pauses (silences longer than 50 ms) which were in turn randomly prolonged or shrunk (*jittered* condition; **Fig 1**, see Methods section for details). This manipulation altered the distribution of pauses, keeping their overall length but increasing the standard deviation (SD = 0.21 for *control* and SD = 0.58 for *jittered;* **Fig 1D**). Thus, we inherently changed the temporal regularities of natural speech between words and syllables [45,46], increasing the temporal unpredictability but keeping intelligibility intact [39]. Previously, this manipulation has been reported (using EEG) to disrupt speech-to-brain entrainment in the delta range [0.5 - 2 Hz; [39]], which we further aimed to disentangle from contextually-driven encoding in the phrasal scale.

**Figure 1.**
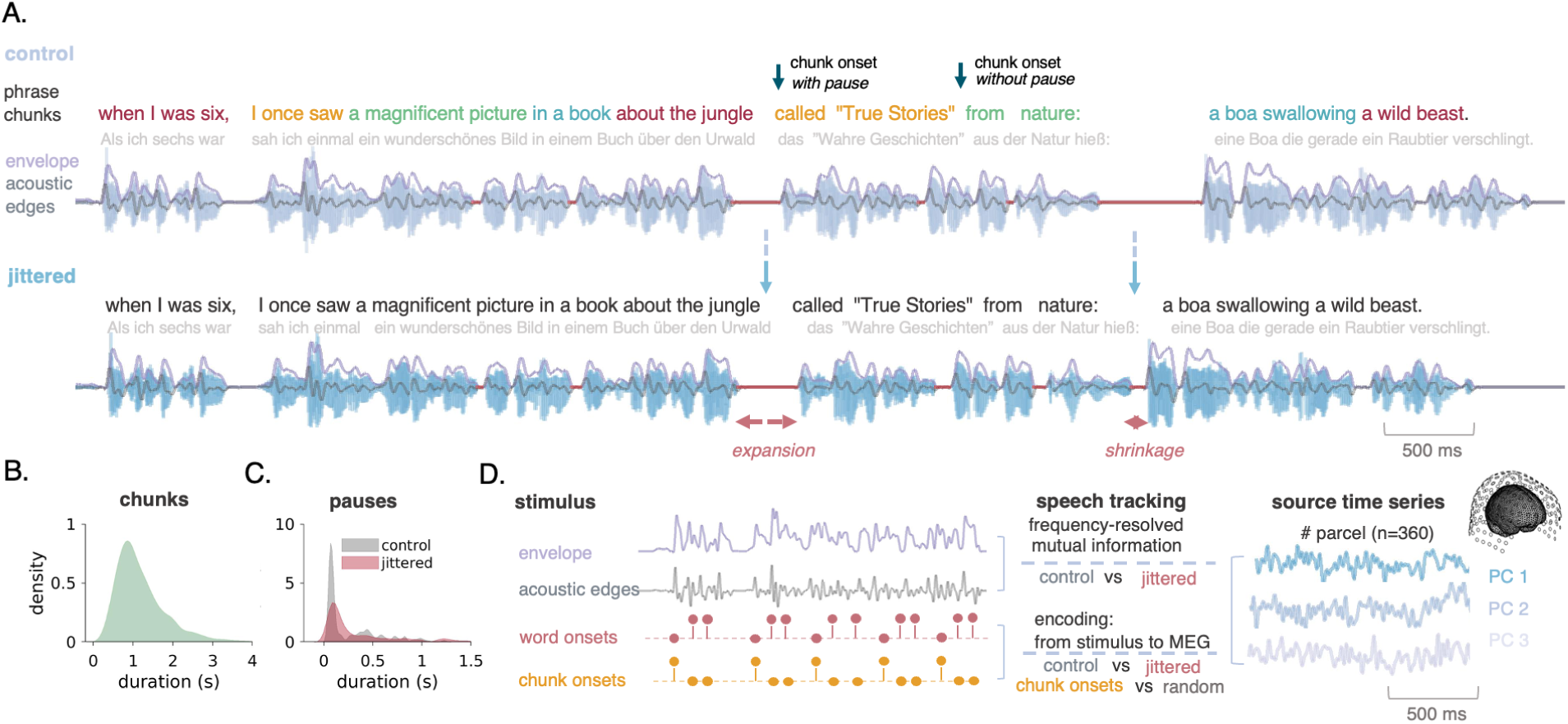
Experimental design and analysis pipeline. **A.** Depiction of the acoustic signal along with the respective phrases (chunks). Text transcriptions were inserted in an language-agnostic morphosyntactic algorithm which automatically identifies chunks from universal dependencies (*chunks* are depicted with different colors on top of original speech signal; [40]; see Methods for details). We identified segments of silence exceeding a duration of 50 ms (red; top) and we randomly increased or decreased their respective length (red; bottom) preserving the mean of pauses duration (see Figure 1C) but increasing the jitter (i.e. the standard deviation) by 90 %, making the temporal structure of speech stream unpredictable (jittered condition). Note that English translation is shown for illustrative purposes only. **Β.** Distribution of chunk duration concatenated for both conditions. **C.** Probability density functions representing pauses length distribution for the two conditions (*top).* The standard deviation was increased after the manipulation, but the overall median was left intact after the manipulation (median = .101; SD = .28 for control, median = .17; SD = .51 for jittered). **D.** The overall pipeline of the study: Acoustic and linguistic features were extracted from the speech waveform (envelope, acoustic edges, word onsets and chunk onsets) along with source time series from the HCP-MMP1 atlas (n = 360; 3 principal components per parcel; [49]). We applied two methods of speech-tracking analysis: First we associated the acoustic features and the source time series with a multivariate frequency-resolved mutual-information analysis for both conditions (see Figure 2), and second we leveraged an encoding’s model approach to predict source time series from linguistic linguistic features, while accounting for the acoustics (see Figure 4).

We first computed the overall amplitude-modulation of the speech signal (envelope) along with its derivative (hereafter we will refer to it as *acoustic edges*) and estimated source activity from 360 brain areas, for both conditions (**Fig 1E**, upper part). We represented the temporal dynamics of each brain area with the first three principal components of the concatenated vertex-level activity within a parcel, as we have observed that this accounts for over 90% of the total variance in auditory areas [47]. Then, we quantified multivariate speech-to-brain alignment [47,48] via frequency-resolved non-linear statistical associations [mutual information (MI); see methods].

The speech-brain coupling analysis resulted in MI values for each area, frequency (from 0.5 - 40 Hz; logarithmically spaced) and time-lags [from -300 to 300 ms; see **Fig 2A** (*upper left)* for representative spectra]. As expected, a significant difference between the two conditions (control vs jittered) was found in the delta frequency range [0.5 - 1.5 Hz; **Fig 2A** (*upper right* and *bottom)*] which was located in bilaterally auditory cortices (group statistics; p<.05; cluster corrected; **Fig 2B)**. Thus, we find that disrupting prosodic punctuation of natural speech compromises bilateral speech-brain coupling in the delta frequency range.

**Figure 2.**
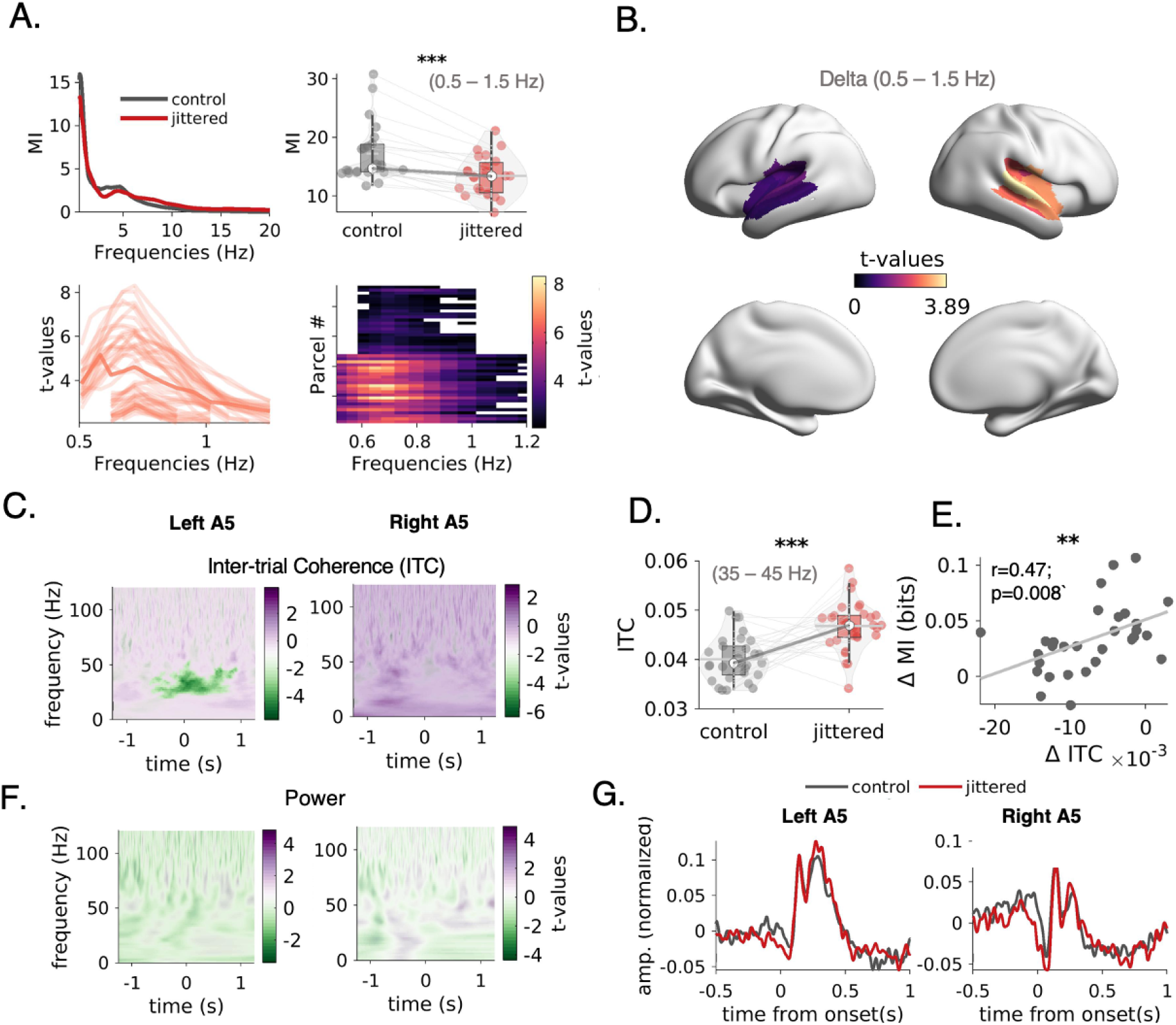
Delta entrainment and gamma coherence are anti-correlated during listening. **A.** Grand average of mutual information (MI) spectra for the control (grey) and jittered (red) condition across significant parcels after a non-parametric cluster permutation test (control vs jittered; *upper left*). Violin-plots depicting the group difference in delta speech brain-coupling (0.5 - 1.5 Hz; *upper right*). Individual T-values spectra from the control vs jittered comparison of MI spectra (cluster-level corrected*; bottom).* **B.** Cortical map of t-values summed across significant frequencies in the delta range (0.5 - 1.5 Hz) **C.** T-values maps from non-parametric cluster permutation test (control vs jittered) of ITC maps for left and right A5. Significant cluster was found in the gamma frequency range (25-45 Hz) centered at speech onsets (group statistics; p<.05; cluster corrected). **D.** Box plots depicting the group difference in gamma ITC between control and jittered condition **E.** Delta speech-brain coupling decrease and gamma coherence increase are correlated (r(29) = .47; p = .008) **F.** T-values maps from non-parametric cluster permutation test (control vs jittered) of power maps for left and right A5. No significant cluster was found (group statistics; p>.05; cluster corrected). **G.** Evoked responses for left and right A5 during speech onsets for control (grey) and jittered (red) condition. No significant difference was observed between the two conditions (non-parametric cluster permutation test; p>.05)

Next, we investigated differences in the temporal dynamics of bilateral auditory cortices (left and right A5) time-locked to speech onsets that follow a predictable (*control*) or unpredictable (*jitter*) pause. First, we focused on phase alignment to the stimulus by means of intertrial coherence (ITC; [50]) for the time range from -1.5 s to 1.5 s relative to speech onset. We identified trials per condition and computed ITC for both conditions (n=506 for control; n=329 for jittered, pseudorandomized across subjects) within bilateral A5 and statistically compared them by means of non-parametric permutation tests (see Methods). We report a left-lateralized increase in gamma ITC (25-45 Hz, p<.05) during a presentation of temporally unpredictable speech onset which was time-locked to speech onsets (**Fig 2C** and **Fig 2D**). Strikingly, the observed increased gamma ITC at the jittered condition was significantly correlated with the decreased delta entrainment across participants (r(29) = .47; p = .008; **Fig 2E**) suggesting a functional relationship between these effects; when prosodic-related delta speech-brain drops, gamma coherence increases. Interestingly, we did not find any significant difference between the frequency-specific power (**Fig 2F**) of the epochs centered on speech onsets or in the evoked responses (p>.05; **Fig 2G**).

Importantly, we have parametrically altered the sensory-driven distributional properties and parsing of incoming speech which we further aimed to evaluate whether it affects sampling in the time-scale of phrases. All in all, our findings suggest that alterations in the temporal structure of speech – which leads to reduced predictability of onsets – disrupts speech-brain coupling in the delta frequency range which is compensated by an increase in gamma coherence during sampling of speech onsets.

### Multi-word chunk onsets elicit delta alignment in bilateral auditory areas

We then asked whether delta alignment in auditory cortices is evident at the boundaries of multi-word chunks. In usage-based syntactic processing models from psycholinguistics and cognitive psychology, these abstract units are thought of as sets of words that depend on each other (e.g., a determiner, adjective, and noun). While memory limits constrain chunk duration and thus necessitate occasional segmentation, each chunk-size filling of the buffer is coherent through the inter-word dependencies—thus allowing for the integration of word meanings within the chunk, and thus comprehension [43]. Previously, invasive work has shown that a spatially distinct area within superior temporal gyrus is exclusively sensitive to speech onsets [51] while non-invasive recordings demonstrated that delta speech-brain coupling is specific – and temporally restricted - to speech onsets [23]. However, it is not yet clear whether contextually-driven onsets (such as the boundaries of multi-word chunks) are sufficient to align activity on longer time-scales during listening. To address this, we leveraged a morphosyntactic and language-agnostic algorithm [40] to annotate phrases as multi-word chunks in a natural speech paradigm (**Fig 1A**; each chunk is depicted with different color). This method accounts for dependency annotations, yields optimal sub-trees in the dependency trees – thus aligns with major syntactic rules [40–42]. When conflicting chunks were identified in the process, optimal ones were selected based on information theory and part-of-speech tags (see Methods). In total, identified chunks [n = 868 (447 for control and 421 for jittered, pseudorandomized across participants)] with a mean length of ∼1 second (mean = 1.21 s; SD = 0.65; **Fig. 1C**). First, we marked segments of speech envelope trials centered on speech onsets and chunk onsets (**Fig 3A**; left). For the later, we wanted to isolate chunk onsets from prosodic boundaries, so we only considered chunk onsets that were not discernible from a preceding pause (pauses exceeding duration of 50 ms; n=112 for control and n=161 for jittered, counterbalanced across participants; **Fig 3B**). This was done to ensure that beginnings of chunks occur without acoustic onsets. It is evident that whereas speech onsets with a pause have a distinct peak in the averaged speech envelope, this is not the case for the onsets without a pause (**Fig 3B**, plots on top).

**Figure 3.**
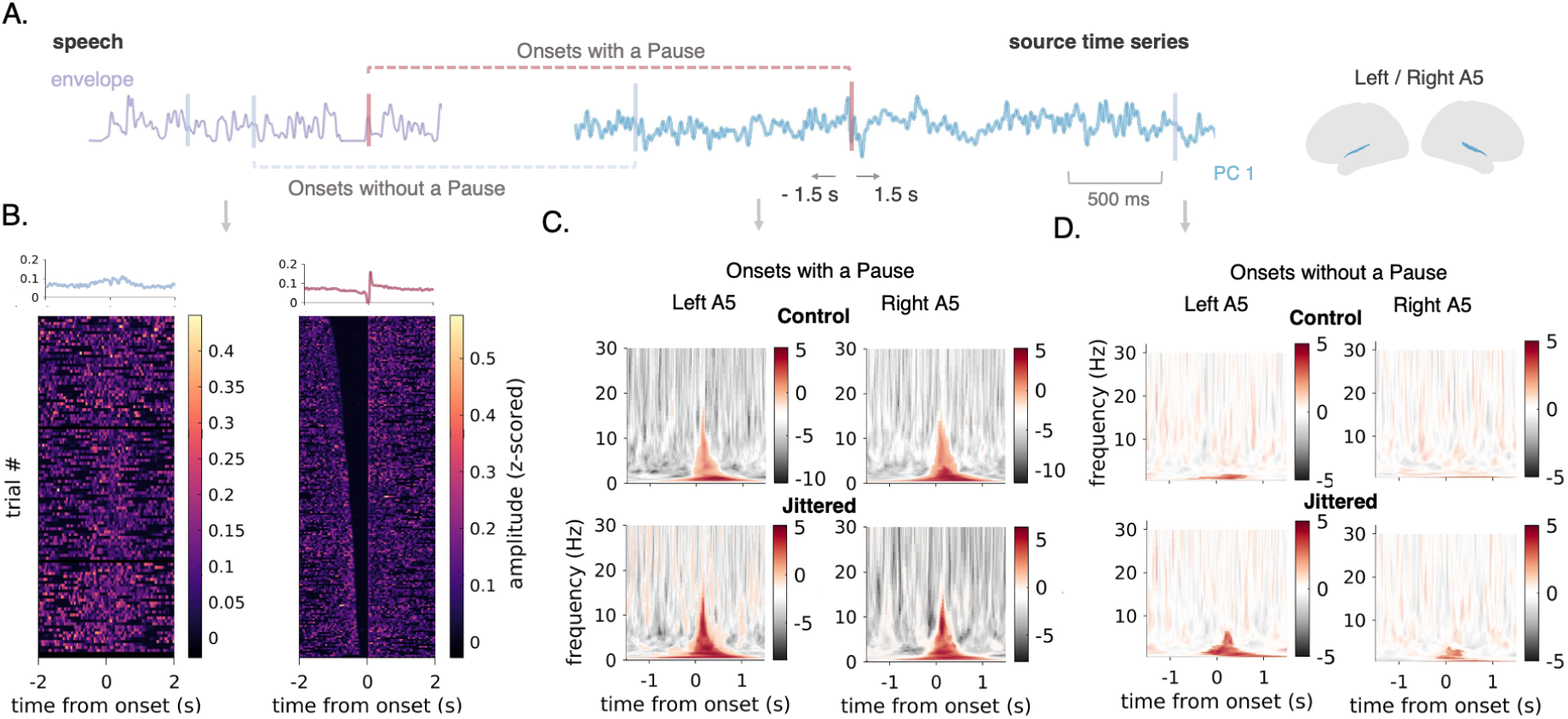
Delta inter-trial coherence elicited by speech and chunk onsets. **A**. Schematic illustration of the analysis: source-derived trials from left and right A5 for onsets with and without a pause. Length of the trials was 3 seconds. **B.** Speech envelope trials for chunk onsets without a pause (*left*) and speech onsets sorted by the preceding pause length (*right*). **C.** T-values of ITC maps between trials of onsets with a pause vs null trials, after nonparametric cluster permutation test for control (upper) and jittered (bottom) condition. Significant clusters are indicated with higher opacity (*red*) **D.** T-values of ITC maps between trials of onsets without a pause vs null trials, after nonparametric cluster permutation test for control (upper) and jittered (bottom) condition. Significant clusters are indicated with higher opacity (*red*).

We then extracted bilateral auditory activity (from left and right A5) during the presentation of these segments using the first three principal components. From the continuous source’s estimated MEG data we extracted frequency-resolved trials using continuous morlet wavelet decomposition (**Fig 3A**; see Methods for details). We computed inter-trial coherence (ITC) as a measure of phase alignment within single trials of bilateral auditory activity and we statistically compared it to null trials (i.e. randomized triggers).

As expected, we found a strong phase consistency in the delta band time-locked to chunk onsets after a pause, extending to theta frequency range for both control and jittered conditions (group statistics; p<.05; cluster corrected; **Fig. 3C**). Strikingly, we also found delta alignment for chunk onsets not distinguished by a preceding pause in the 1st principal component (group statistics; p<.05; cluster corrected; **Fig 3D**). Similarly, delta alignment is also evident in the 2nd and 3rd principal components (**Supp. Fig 1A)**. Thus we report that delta alignment is evident during acoustically-driven speech onsets and independently by contextually-driven chunk onsets which were not marked by an acoustic boundary.

### Contextually-driven encoding of multi-word chunks in bilateral auditory and frontal areas

Next, we wanted to assess whether including information of chunks in encoding models improves the predictive power of neural responses. This would serve as further evidence for contextually-dependent processing of speech, independent from distributional properties of acoustics. To this end, we used a time-resolved temporal response function (TRF) approach [52,53] to model neurophysiological fluctuations of the source-estimated signal (n=360 parcels; separately for the first three principal components) from a set of regressors. Specifically we estimated regression-weights between neurophysiological responses and the speech envelope, acoustic edges, word and chunk onsets for different time-lags (-100 to 1 s; in steps of 10 ms). This analysis first aimed to pinpoint the predictive power of chunks above-and-beyond word level processing and acoustics. To this end, source-reconstructed brain activity was modeled with ridge regression in a train data set from a set of acoustic (speech envelope and acoustic edges) and linguistic features (discrete word and chunk onsets).

In **Fig. 4A** we plot the estimated regression weights within left and right A5, for the first principal component. We find a typical temporal response profile for the speech envelope and the acoustic edges (bilateral peaks at 60/200 ms and trough at 130 ms), matching a damped wave in the theta frequency range, and thus reflecting the rhythmicity of speech [54]. Note that this profile is also captured from the 2nd and 3rd principal component (**Supp. Fig 2A)**. In contrast, the temporal response profile of word and chunk onsets depict a response in a slower frequency range. Specifically, word onsets show peaks at 90 and through at 220 ms, which are observable only in the 2nd principle component **(Supp. Fig 2A)**. Strikingly, the regression weights for the chunk onsets show a partly similar but mostly distinct profile from word onsets across the three principal components. We see distinct peaks (at 140 and 290 ms) for the left and at 120 ms for the right, with bilateral troughs around 520 ms. This difference indicates that chunk onsets might have predictive power above-and-beyond word level information.

**Figure 4.**
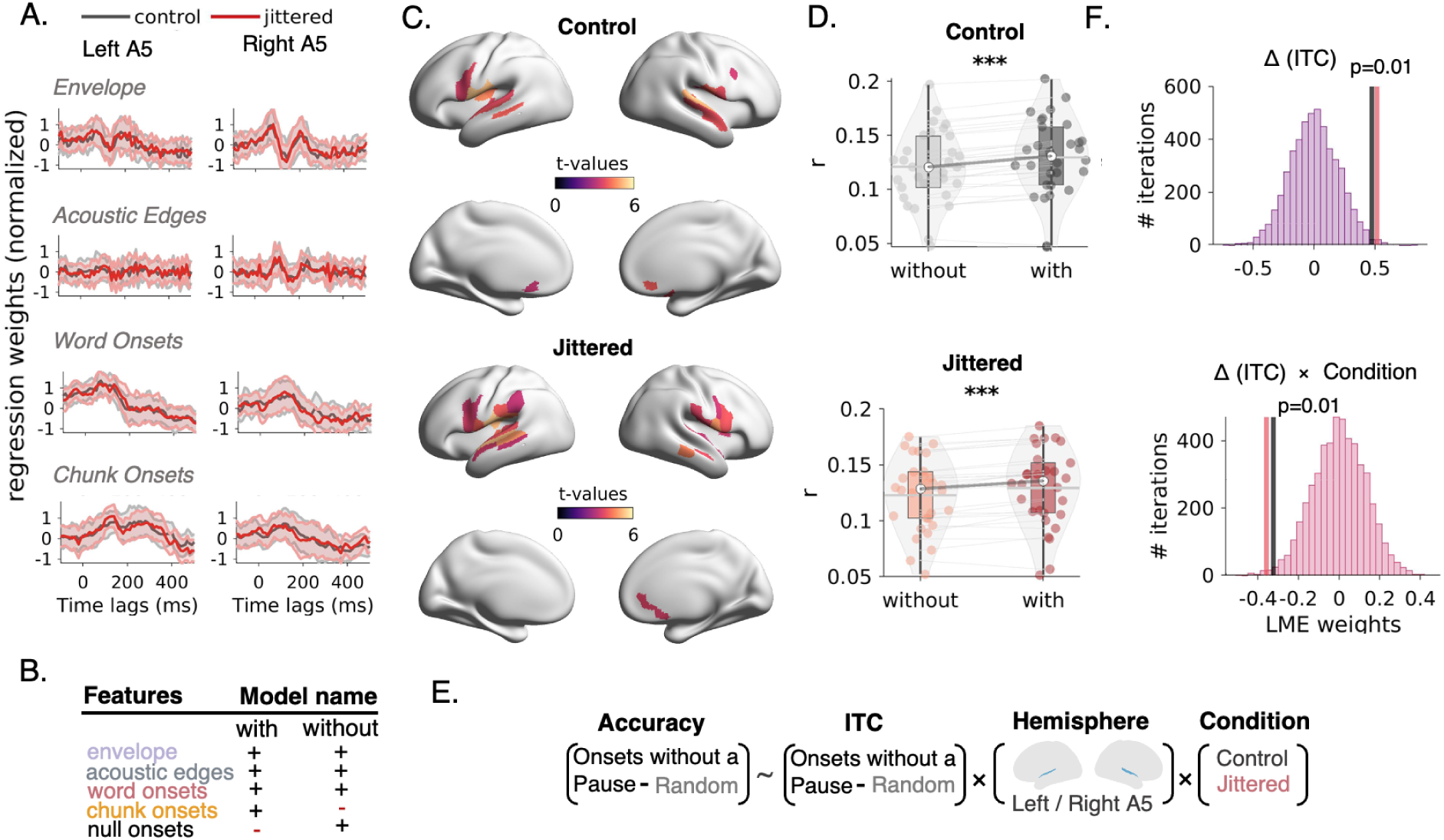
Contextual tracking of chunks in bilateral auditory and frontal areas. **A.** Temporal response functions across feature spaces (envelope, acoustic edges, word onsets, chunk onsets) for control (grey) and jittered (red) conditions **B.** Feature spaces used for the two models: In one model the envelope, acoustic edges, word onsets and chunk onsets were used (with) and in the other model the chunk onsets were replaced by null pulse responses (without) **C.** Pearson’s correlations of predicted responses per parcel (n=360) and principal components (n=3) were averaged across brain areas and subjected into non-parametric permutation cluster test to the performance of the two models (with vs without pauses). Cortical maps of t-values of the cluster t-test for control and jittered condition **D.** Violin-plots depict the difference in accuracy (as expressed by Pearson’s correlation) between the models with and without chunk onsets, extracted and summed across significant parcels **E.** Schematic illustration of the linear mixed effect model (LMEM) employed: Increase of accuracy between the two models (with and without) was expressed as a linear combination, along with their interactions, of ITC increase in chunk onsets without pauses, from left and right A5, for control and jittered conditions **F.** Histograms shows the empirical LMEM weight for the [Δ (ITC); top] and Δ (ITC) x Condition interaction against the null distribution computed from 5000 random iterations of subjects’ accuracy vectors.

To test this hypothesis, we compared the cross-validated performance between predicted and empirical time-series by means of Pearson’s correlation using two models: One model encompassing all predictors (envelope, acoustic edges, word onsets and chunk onsets; hereafter we will refer to that as *full* model) with another model including the same predictors, except that the chunk onsets were jittered randomly (null onsets; we will refer to that as the *no chunks* model; **Fig 4B**). We computed the correlation between the predicted and empirical time-series for each parcel and each principal component and averaged across components. Then, we compared the performance of the two models by means of cluster-based permutation testing. We found a striking difference between the two models for both conditions located in bilateral auditory, expanding to frontal areas (group statistics; p<.05; cluster corrected; **Fig 4C**) in which the model with the chunk onsets performs significantly better compared to the model without chunk onsets (**Fig 4D**). As expected from the lack of difference in regression weights between conditions, there was no significant interaction between the construction accuracy of the two models (with and without) and conditions (control and jittered; group statistics; p>.05)

Next, we asked whether the observed increase in performance can be predicted by the magnitude of delta alignment at chunk onsets without pauses (see **Fig 3D**). For this we employed a linear mixed effect model (LMEM) approach in which we expressed the difference in accuracy between the two models (with and without pauses) by the linear combination of the delta phase concentration (as depicted in ITC values) of left and right A5 for control and jittered condition; along with their respective interactions (see **Fig 4E** for graphical illustration of the model). This would further serve as evidence that the contextual processing of chunks can be traced down to the phase alignment of delta activity. We found a significant main effect of ITC [t(112)=2.74, p=.007], ITC ✕ condition interaction [t(112)= -3.5, p=.004] and ITC ✕ hemisphere interaction [t(112)= -0.26, p=.03]. To further test the reliability of the model we additionally randomly permuted participants k=5000 times, recomputed LMEM and constructed a null distribution of LMEM weights. Empirical LMEM weight scored above 99th percentile for the ITC (**Fig 4F**, top) and below the 1st percentile for the ITC ✕ condition interaction (**Fig 4F**, bottom). This further corroborates the overall influence of delta alignment on the processing of multi-word chunks.

To summarize, we report an increase in accuracy of encoding models when phrasal onsets information is employed. This increase is located in bilateral auditory and frontal areas, suggesting top-down contextual processing of chunks. Importantly, we find that for the bilateral auditory area A5 this increase can be traced down to the observed delta alignment during chunk (or multi-word) onsets.

## Discussion

How context-invoked processes interact with bottom-up sensory evidence during speech perception remains a fundamental question, for which data has been highly contentious. The main focus of this study was to disentangle top-down contextual processing from bottom-up prosodic features during listening to a story. We find compelling evidence that both these processes co-exist independently and can be both traced down interactively in the phase of slow rhythmic cortical activity in the delta frequency range. Our results have important implications for the interaction of on-line sensory evidence and long-term contextual knowledge during spoken language processing.

We confirmed that temporally manipulating prosodic cues compromises cortical alignment to speech, specifically in bilateral auditory areas in the delta frequency band (**Fig 2A-B**). This replicates previous EEG findings [39], aligns with a proposed role of delta oscillations in temporal expectations [55,56] and is consistent with previous speech-brain coupling research proposing delta activity specific to speech onsets [23]. In this line, we foster the notion that delta speech-brain coupling reflects top-down predictive processes [15] evoked by acoustical regularities [57,58]; in particular the temporal structure of pauses [46]. While processing temporally unpredictable speech-onsets, we moreover report an increase in gamma synchrony, as expressed by higher inter-trial coherence (**Fig 2C-D**). Auditory-evoked gamma band responses (GBR) have been previously reported and dissociated from the slow-frequency evoked component [59]. Broadly, gamma oscillations have been linked to sensory prediction errors realized – on the microcircuit level – by collective firing of superficial pyramidal neurons [60,61]. Here, these might serve as a speech-specific [62] feedforward index of temporal prediction-violation [63] and consequently rhythmic gain modulation [64] across the auditory pathway hierarchy, as previously shown in the visual domain [65,66]. Interestingly, we find that these two processes are anticorrelated: The larger the increase of gamma synchrony, the smaller the decrease of delta speech-brain coupling observed across participants (**Fig 2E**). Although this correlation does not serve as causal evidence, it suggests that delta speech tracking and gamma synchrony in speech onsets subserve feedback and feedforward operations respectively, such that when feedback predictions are thwarted, feedforward projections counteract during processing speech onsets.

Moreover, we find no evidence of power difference between temporally predictable and unpredictable conditions in the alpha frequency band within bilateral auditory areas (**Fig 2F**). This suggests that the previously reported alpha band power effect is specific in frontal and higher-level associated areas [39] which can further interact with speech-specific networks [67] and support language processing [28,68]. Importantly, we showed that temporally disrupting prosodic boundaries in speech alters a feedforward and feedback processing balance of temporal tracking during listening.

As expected, we found strong delta alignment during speech onsets marked with a prosodic boundary [22] in auditory areas, irrespective of the temporal predictability in the speech signal (**Fig 3C**). This points to the direction that perception of prosodic boundaries through intrinsic delta-band activity [20] constitutes sensory-driven bottom-up process and leads to on-line segmentation of speech [69,70]. Prosodic boundaries are also evident as a closure positive shift in the event-related component [71], but whether these two phenomena are inter-related remains an open question. Nevertheless, our results support the notion that acoustically-related processing of prosodic phrases is occurring in a bottom-up fashion through a delta-band frequency channel. This points to a more complex pattern than previously hypothesized top-down and bottom-up processing via low- and high-frequency channels respectively [66,72], as previously reported in the visual [73] and auditory domain [74].

Delta alignment was also evident in bilateral auditory areas at chunk onsets, not marked by a prosodic boundary (**Fig 3D**). Naturally, no difference was observed between the two conditions as the chosen speech trials had not been manipulated. We consider this intrinsic auditory delta alignment as a manifestation of top-down or contextually-driven cortical activity during listening to naturalistic speech. Previously, using an encoding model approach, it was shown that sentence context – extending beyond entropy, surprisal and sensory distributional facets – modulates word processing in the amplitude of the delta band [75]. Here, we show that this modulation can be established through processing of computationally-formulized speech chunks, derived from one-level dependency treebanks. Moreover, increased speech-brain coupling has been reported, as expressed by higher mutual information values, between the phase of the delta band and pulse-train-coded phrasal chunks when compared to jabberwocky and prosody-controlled speech chunks [76,77]. We extend these findings in naturalistic listening by leveraging a combination of the encoding model approach with linear mixed modeling: We show that the increase in model accuracy from chunk onsets information can be predicted by the instantaneous delta activity in bilateral auditory areas. Again, this does not constitute a causal proof of interaction, but it strongly suggests a functional connection between chunk processing and the phase of delta activity in bilateral auditory areas.

Chunks onset information drastically increased the accuracy of encoding predictive models in temporal and frontal areas, for both control and jittered experimental conditions. Before, similar encoding models including only acoustic information of the signal (envelope and derivative) were superior (in terms of accuracy) to ones including more sophisticated linguistic-related information (articulatory features; [78]; but see [79]). Here, the profound increase in accuracy with chunk onsets might reflect long-term memory processes which align (in a feedback manner) to contextual dependencies acquired through development [80], and facilitating predictions [81], accessed through long temporal integration windows [82].

Consequently, our findings echo the analysis by synthesis (A × S) framework of speech processing [83,84], within which speech input is processed by concurrently matching internally-generated brain states (synthesis; feedback) to the incoming acoustical signal (analysis; feedforward). In this line, lexical representations are subject to feedback contextual inferences, prior to sensory input [85]. We propose that in the phrasal timescale this might be operationalized at the neural level by delta-band alignment. As proof-of-principle, delta-band entrainment (1.6 - 1.8 Hz) was evident for several cycles beyond the sensory stimulation [57], while persistent entrainment beyond stimulation was found to affect perceptual inference during listening [86].

Furthermore, we found no difference in the increase of encoding accuracies between the two experimental conditions (control and jittered). This might reflect the fact that for both conditions the speech envelope and acoustic edges were controlled. We note that reduction of speech-brain coupling in the delta frequency band was observed in the phase (not in the power) of bilateral auditory activity; the encoding models operating in the power of the broadband signal remain blind in this effect. Future studies could implement such an encoding approach, utilizing the phase of cortical activity as a dependent variable. Importantly, the interaction between condition and intertrial coherence in the delta frequency band (**Fig 4F**, bottom) indicates that delta alignment at chunk onsets serves as a better predictor of the model, leveraging chunk onsets information, in the jittered compared to the control condition. While surprising, as for both experimental conditions chunk onsets without pauses were considered (and thus there was no prior manipulation), this probably stems from the slightly higher (but statistically insignificant) ITC values in the jittered condition (**Fig 3C,D)**.

## Conclusion

In summary, we show neural evidence for parallel processing of prosodic and contextual sampling during listening to a story. Contextually-driven processing is underpinned by the alignment of slow cortical activity (in the delta frequency band), which matches the phrasal timescale, and it differentiates from the speech-brain coupling in this frequency range: delta speech-brain coupling seems to be under predictive top-down control, while intrinsic delta alignment acts in a bottom-up fashion during processing of speech onsets. All in all, we find a temporally and functionally dissociated role of intrinsic and bilateral auditory alignment of dynamics within the delta frequency band, one for contextual-driven and another for prosodic sampling of speech.

## Methods

### Participants

Thirty native German speakers (19 females; mean age = 25.21 years, age range 18 - 31 years) participated in this study. All participants provided written consent prior to the experiment, were instructed explicitly with written instructions and received a monetary compensation of 15€ per hour. The study was approved by the Ethics committee of the Medical Faculty of University of Münster (application number: 2018-066-f-S) and was conducted according to the Declaration of Helsinki.

### Stimulus and study design

Participants listened to a German translation of the story *Le Petit Prince (*The Little Prince) extracted from a large multilingual corpus for research (five chapters; total length = 19.31 minutes) narrated by a native female speaker. We presented isochronous original and manipulated (jittered) recordings of the story. For the latter, pauses of speech were identified and their overall pause duration distribution was jittered, preserving the mean but increasing the standard deviation by randomly shortening or lengthening each pause, using custom Matlab scripts from [39]. Specifically, to detect pauses we computed the wide-band amplitude envelope of the speech signal by band-pass filtering into 11 logarithmically spaced bands between 200 and 6000 Hz and averaged the narrow-band Hilbert-transformed envelopes. The wideband envelope was further smoothed with a 10 ms gaussian filter and normalized. Periods in which amplitude was < 0.1 were considered as silence and when a silence was exceeding at least 50 ms duration we considered it as a pause. For each pause we systematically increased or decreased the length, increasing the jitter (i.e the SD of the pause distribution) by 90%. We reconstructed the speech signal with the manipulated pauses assuming zero amplitude during each pause and we cosine ramped for 5 ms the onset and offset of speech material around each pause. To mitigate minor acoustic artifacts, a continuous white noise background with relative RMS level of 0.05 was added to the original speech signal.

Speech material was delivered in two experimental blocks: One block included the first two chapters and the second the rest of the three (total duration = 9.73 and 9.58 minutes). The order of experimental conditions and blocks were pseudorandomized across subjects. To assess whether participants paid attention to the story, they answered by means of a button press 15 multiple choice questions (3 after each chapter; 2 response options each).

### Identification of multi-word chunks

Multi-word chunks were extracted from the text using a morphosyntactic and language-agnostic evolutionary algorithm which identified chunks from universal dependency (UD) treebanks [40]. We applied the chunking algorithm from [41]. The text was parsed automatically obtain UD annotations by using the English Web Treebank model from UDPipe v2.6, which has a high performance with a labeled attachment score of 87.43% on the raw text of the corresponding test file [87,88]. We note that chunks were defined as words and bound morpheme sequences with all possible local dependency clusters [40,42]. Initially, *chunks* were considered as base-level subtrees. Then, unitary chunks were minimized with the following procedure:

For a given sentence and its associated tree structure, potential multi-word chunks are extracted if, for the node n at the position x with the corresponding head ℎ at position 𝑘 (where k can be either greater or less than x), the nodes between x and k-1 (if k>x) or the between x and k+1 (k<x) have the same head h. This process results in overlapping chunks, where some words of a chunk might be a dependent of another chunk. To select optimal chunks for a given tree, we computed normalized pointwise mutual information (NPMl; [89])) between the Universal part-of-speech (UPOS) tag of a node (t) and the tuple of the UPOS of the head of that node (ht) and the relation between the node and its head (rel) for a given node, as described in (1) and (2). Then we average NPMIs associated with the nodes within a candidate chunk (see (3)). Chunks with higher average NPMI are selected. Subsequently, to minimize unitary chunks (i.e., with only one node), we removed punctuations with the UPOS tag of punctuation only and then attached remaining floating unitary nodes to the chunks where those unitary chunks are syntactically linked to any element of the corresponding chunk.

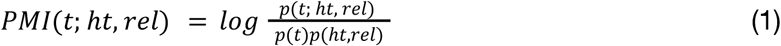

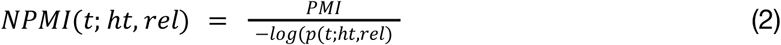

with the average NPMl for a candidate chunk defined as:

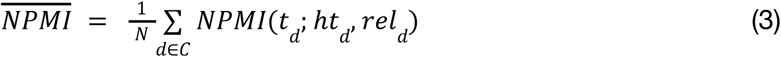

where 𝑁 is the total number of nodes in a phrase and 𝑑 is a dependent in a given phrase 𝐶. We further refer the reader for a more detailed description of the methods used in [41].

### Data acquisition

Evoked magnetic fields were recorded in a magnetically shielded room via a 275 channel whole-head MEG system (OMEGA, CTF Systems Inc, Port Coquitlam, Canada). Data were continuously recorded with a sampling rate of 1200 Hz. Subjects were seated upright with a fixed head position, comfortably stabilized inside the MEG dewar via pads. The stimulus was delivered using PsychToolBox [90] with two 60 cm long silicon tubes. Questions after each chapter were projected onto the back of a semi-transparent screen positioned approximately 90 cm in front of the subjects’ nasion using an PROPixx Lite Projector (VPixx Technologies Inc., Canada) with a refresh rate of 60 Hz.

### Data preprocessing

Data were preprocessed with the MNE-python toolbox [91]. Initially, we epoched the data according to the onset and offset of each chapter. Then, we filtered the data with a Hamming-windowed finite impulse response (FIR) zero-phase high-pass filter with a 1 Hz cut-off. We extracted 40 independent components for the 2nd-order gradiometer data using the *fastica* algorithm. We manually identified artifactual components belonging to eye movements and heart activity (mean number = 4.75, SD = 1.58) and we reconstructed the original recording in which the artifactual components were removed. We filtered above 1 Hz prior to the ICA to avoid biases towards lower frequencies, which tend to have greater power [92]. Finally, we filtered the reconstructed data with a Hamming-windowed FIR zero-phase filter filter with a 0.2 Hz cut off and we downsampled the data to a sampling rate of 600 Hz.

### Source localization

T1-weighted Magnetic Resonance Images were obtained from each participant in a 3-T scanner (Gyroscan Intera T30, Philips, Amsterdam, Netherlands). 400 contiguous T1-weighted slices of 0.5 mm thickness in the sagittal plane (TR = 7.33.64 ms, TE = 3.31 ms) were collected by a Turbo Field Echo acquisition protocol. The field of view was set to 300 ✕ 300 mm with an in-plane matrix of 512 ✕ 512 setting defining the native voxel size at 0.58 ✕ 0.58 ✕ 0.58 mm^3^. The intensity bias of the images was then regularized using SPM8 (Statistical Parametric Mapping, in order to account for intensity differences within each tissue and the images were resliced to isotropic voxels of 1.17 ✕ 1.17 ✕ 1.17 mm.

Individual T1-weighted MRIs were coregistered to the MEG coordinate system, aligned with the digitized head shapes using custom-made markers in participants’ earmolds during MR image acquisition using the fieldtrip toolbox for MATLAB 2021b [93] (MathWorks Inc.). To describe an individual subject’s cortical sheet, we performed cortical surface reconstruction with the Freesurfer image analysis suite, using the surface-based stream implemented in the recon_all command-line tool. The post-processing of the reconstructed cortical surfaces was performed using the Connectome Workbench wb_command (v1.1.1; https://www.humanconnectome.org/software/workbench-command). The cortical sheet reconstruction procedure resulted in a description of individual subjects’ locations of potential neural sources along the cortical sheet (source model) with 64984 source locations (vertices) per hemisphere. We generated a single-shell spherical volume conduction model (head model) based on a realistic shaped surface of the inside of the skull [94] and computed the forward projection matrices (leadfields).

Source activity was estimated by computing linear constrained minimum variance (LCMV) beamformer coefficients from the MEG time-series for each vertex on the cortical sheet [95]. The sensor covariance matrix used was computed across all trials. The lambda regularization parameter was set to 5% and time series were extracted for each dipole orientation, resulting in three time-series per neural source. To reduce the dimensionality of the data, we applied an atlas-based parcellation of cortical space, resulting in 180 ROIs per hemisphere [96] and applied principal component analysis within a parcel after concatenating time-series from each vertex and orientation. Then, we extracted the first three principal components, as they account for more than 90% of the total variance in auditory areas [47].

### Spectral estimation

To identify frequency-specific activity for both conditions (control and jittered), we applied a continuous wavelet transformation in each parcel time-series (CWT; cwtfilterbank.m and wt.m) for 80 logarithmically spaced frequencies (from 0.1 Hz to 120 Hz) using the analytic Morlet wavelet with the symmetry parameter (γ=3) and time-bandwidth (tb=10). For more accurate depiction of the continuous signal, a L1-normalization is implemented in the CWT algorithm, assigning equal magnitude to components with equal amplitude across different scales. With this we obtained a complex-valued continuous time series for 360 parcels, 3 principal components and 80 frequencies, from which we extracted phase angles and amplitude from epochs ranging from -1.5 to 1.5 from gap or chunk onsets. Post-onset amplitude was further normalized by subtracting the mean and dividing by the standard deviation of pre-onset values (-1 s to onset).

#### Inter-trial coherence (ITC)

We estimated ITC from summing all the phase angles from epochs ranging from -1.5 to 1.5 s from the onset of a trial, according to the following formula:

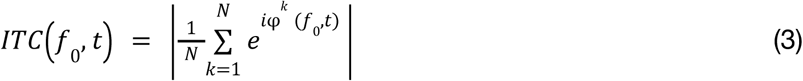

Where N is the number of trials, *φk* is the local phase angle of the signal at the current trial (in radians) and 𝑓_0_ the frequency.

### Speech-tracking

To quantify dependencies between the estimated source activity and speech we employed two different approaches: first we computing frequency-resolved speech-brain associations using mutual information and then time domain predictions from stimulus to brain using encoding models. Note that for this analysis source-space data were further downsampled to 100 Hz to reduce computational resources.

#### Frequency-specific mutual information

Initially, statistical dependencies between the speech envelope along with the first derivative [97] and the source space parcels were computed on the basis of information theory [98]. We estimated multivariate gaussian copula mutual information (GCMI) between frequency-resolved multivariate speech signals and source time-series [97]. For the frequency analysis we followed the spectral estimation pipeline described above but restricted to 64 frequencies (from 0.1 to 40 Hz). Following previous work [7,39] we estimated GCMI on the real and imaginary part of the signal as a 2D vector, after normalizing with the frequency-specific amplitude. We estimated GCMI for temporally shifted versions of the stimulus with respect to brain signal for -300 to 300 ms with a 10ms step (total of 61 delays). Additionally we randomly shifted 200 times the speech signal with respect to the source time-series [99] via a circular wrapping around the edges and we generated a distribution of 200 surrogate MI values for each frequency, delay, and parcel. From the computed GCMI we subtracted the mean and divided by the standard deviation of this surrogate distribution and we obtained normalized GCMI values for each frequency, parcel and delay.

#### Forward or encoding models

We used ridge regression to model brain responses from a set of acoustic (envelope and envelope’s derivative) and linguistic features (word and chunk onsets), assuming that neural responses in the source level can be expressed as a linear combination of the these features at multiple time lags [6,100]. This intuitively computes a filter which after a linear convolution with the stimulus feature maps into the brain responses (encoding or forward models).

Specifically, the within-subject linear model we employed can be depicted as:

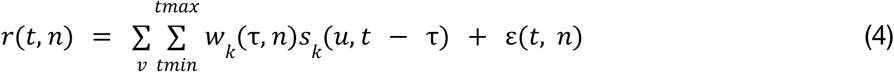

where, the instantaneous brain responses in source level 𝑟(𝑡, 𝑛) of each time sample 𝑡 at each parcel 𝑛 was expressed by the linear convolution of stimulus features 𝑠, with a filter weight 𝑤 across stimulus dimensions 𝑣 and time lags ranging from 𝑡𝑚𝑖𝑛 to 𝑡𝑚𝑎𝑥. Further, ε(𝑡, 𝑛) is a random white noise process, capturing part of the signal which is unrelated to stimulus.

To compute the filter weights we minimized the mean-squared error between source-level responses, 𝑟(𝑡, 𝑛) and the predicted 𝑟(𝑡, 𝑛) as follows:

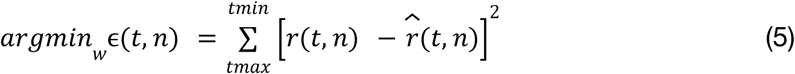

By using the following the closed-form solution of:

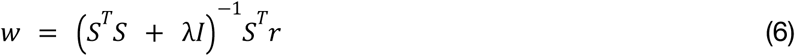

Where 𝑆 denotes the lagged time series of the stimulus representation 𝑠_𝑘_, 𝐼 the identity matrix and λ a regularization term. To leverage the fact that 𝑆^𝑇^𝑆 is unstable, when columns of 𝑆 are correlated and to avoid overfitting of high-frequency noise, a range of regularization terms were tested [lambda values: (10^-6^ - 10^6^)]. We tuned λ using a “leave-one-out” cross-validation framework maximizing the Pearson’s correlation between the observed and predicted responses [101].

Analysis was done using the mTRF toolbox [53]. Specifically, we adjusted λ optimizing the mapping from stimulus-to-brain using *mTRFcrossval,* estimated the filter weights for each stimulus feature using the *mTRFtrain* with 𝑡𝑚𝑖𝑛 = -0.15 s and 𝑡𝑚𝑎𝑥 = 1 s and tested the corresponding model with the *mTRFpredict*. Specifically, Pearson’s *r* was computed between the observed and the reconstructed responses of the left-out trial per parcel (n=360) and principal components (n=3). We note that *r* for each parcel was averaged across the three reconstructed principal components of source activity.

Subsequently, performance of two models were evaluated: one model including the envelope, the envelope’s derivative, word onsets and chunks onsets as stimulus features (we refer to as: *with* chunk onsets) and an identical model in which the chunk onsets were shifted randomly in time (we refer to as: *without* chunk onsets). Even though neural responses can be affected by other predictors in the lexical [word frequency/surprisal [102]] or sub-lexical abstraction [phoneme onsets [52,79]] we decided not include them in the model to leverage the advantage of keeping a minimum number predictors while assuming that the potential unexplained variance by the these factors would be equally modeled through the residuals ε(𝑡, 𝑛) of both models.

### Linear-mixed effect modeling

We used LMEM to model the encoding model’s accuracy (as a response variable) as a linear-combination of fixed effects shared across participants (inter-trial coherence (ITC) of left and right A5 for control and jittered condition, along with their corresponding interactions) and participant-specific random effects. Specifically we set up the following LMEM, using the equation:

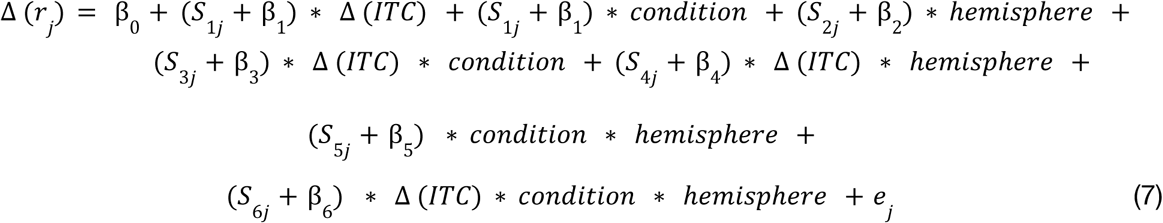

For participant j, the accuracy difference between encoding models with and without chunk onsets was expressed as a combination of an intercept (β_0_), the fixed effects of dependent factors (β_1_… β_6_), random slopes (𝑆_1_j… 𝑆_6*j*_) and an error term (*e_j_* ∼ *N*(0,σ^2^). Random slopes were included in the model to account for intersubject variability, by means that the fixed effects would not modulate the encoding model’s accuracy identically across participants. Furthermore, significance of the empirical LMEM weights was further tested by computing the LMEM n=5000 times after shuffling participant’s model accuracies and comparing its rank with percentiles of the null distribution (1st for negative and 99th for positive values).

### Statistical analysis

Significance in the group level was determined with cluster-based permutation tests [103] using *ft_freqstatistics* in Fieldtrip. For this, a series of 2-tailed *t* tests of individual data points (MI in each frequency, Pearson’s *r,* ITC or amplitude maps) were conducted for each parcel between the two experimental conditions (control, jittered) and resulting t-values were thresholded at p=0.05. Then, spatially and/or spectrally adjacent significant data points were defined as clusters with an assigned cluster-level statistic constituting the sum of the t-values within each cluster. Further, we used Monte Carlo approximation to test each cluster for significance. For that, single subject data points between the two conditions were randomly interchanged and the series of *t* tests, clustering and estimation of cluster-level statistics were recomputed. After repeating this procedure 5000 times the original cluster-level statistics were compared with the histogram of the randomized null statistics. When initial clusters yielded a larger cluster-level statistic than the 95% of the randomized data they were considered as significant.

## Acknowledgments

We acknowledge support by the Interdisciplinary Center for Clinical Research (IZKF) of the medical faculty of Münster (grant number Gro3/001/19). JG was further supported by the DFG (GR 2024/5-1) and RN was supported by a grant from the German Science Foundation (CRC 1451/A07).

