## Supplementary material for "Dissociating endogenous and exogenous delta activity during natural speech comprehension": Supp. Fig.

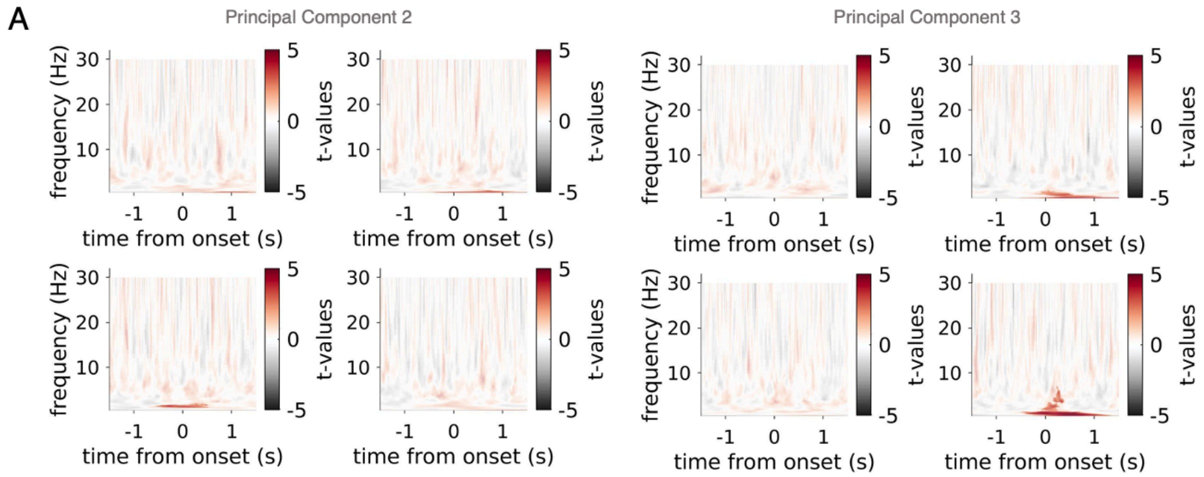


**Supplementary Figure 2**. **A.** T-values of ITC maps between trials of onsets without a pause vs null trials, after nonparametric cluster permutation test for control (upper) and jittered (bottom) condition. Significant clusters are indicated with higher opacity (*red*).

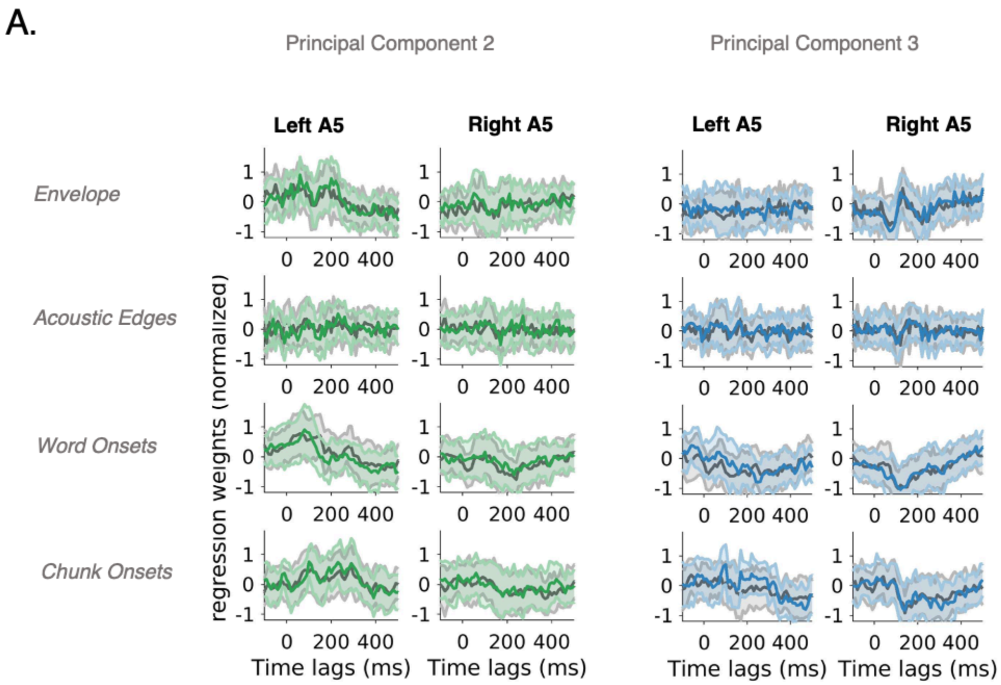


**Supplementary Figure 2.** Temporal response functions across feature spaces (envelope, acoustic edges, word onsets, chunk onsets) for control (grey) and jittered (red) conditions of the 2nd and 3rd principal component.
